# Boreal forests will be more severely affected by projected anthropogenic climate forcing than mixedwood and northern hardwood forests in eastern Canada

**DOI:** 10.1101/2020.11.05.370049

**Authors:** Yan Boulanger, Jesus Pascual Puigdevall

**Affiliations:** Natural Resources Canada, Canadian Forest Service, Laurentian Forestry Centre, Québec, QC, Canada

**Keywords:** Climate change, Harvest, Landis-II, boreal forests, mixedwood, northern hardwood, eastern Canada

## Abstract

**Context:** Increased anthropogenic climate forcing is projected to have tremendous impacts on global forest ecosystems, with northern biomes being more at risk.

**Objectives:** To model the impacts of harvest and increased anthropogenic climate forcing on eastern Canada’s forest landscapes and to assess the strong spatial heterogeneity in the severity, the nature and direction of the impacts expected within northern forest regions.

**Methods:** We used LANDIS-II to project species-specific aboveground biomass (AGB) between 2020 and 2150 under three climate (baseline, RCP 4.5 and RCP 8.5) and two harvest (baseline harvest, no harvest) scenarios within four forest regions (boreal west, boreal east, mixedwood and northern hardwood).

**Results:** Climate change impacts within the boreal forest regions would mainly result from increases in wildfires activity which will strongly alter total AGB. In the mixedwood and northern hardwood, changes will be less important and will result from climate-induced growth constraints that will alter species composition towards more thermophilous species. Climate-induced impacts were much more important and swifter under RCP 8.5 after 2080 suggesting that eastern Canada’s forests might cross important tipping points under strong anthropogenic climate forcing.

**Conclusions:** Boreal forest regions will be much less resilient than mixedwood or northern hardwoods to the projected changes in climate regimes. Current harvest strategies will interact with anthropogenic climate forcing to further modify forest landscapes, notably by accelerating thermophilous species AGB gain in southernmost regions. Major changes to harvest practices are strongly needed to preserve the long-term sustainability of wood supply in eastern Canada. Adaptation strategies should be region-specific.

## Introduction

Increased anthropogenic climate forcing is projected to have tremendous impacts on global forest ecosystems (Heyder et al. 2011). Changes in temperature and precipitation regimes since the last decades have already modified natural disturbances as well as forest productivity in several forest regions worldwide. Among all forest biomes, northern forests will probably be among the most impacted by climate change, with temperatures projected to increase at much higher rates than elsewhere (Price et al. 2013). In Canada, temperatures have already risen by 1.7 °C since 1948, a rate twice as fast as the global average (NRCan 2019). During this time period, wildfire activity has significantly increased (Hanes et al. 2019), boreal tree species productivity have declined (Girardin et al. 2016) while warmer temperatures might be responsible for recent modifications in forest insect outbreak regimes (Pureswaran et al. 2015). Tipping points are likely to be crossed under aggressive anthropogenic climate forcing (e.g., RCP 8.5) potentially resulting in biome shifts (Stralberg et al. 2018). No doubt that these alterations of northern forest ecosystems will severely impact ecosystem services including carbon storage (Kurz et al. 2008), timber supply (Gauthier et al. 2015; Boucher et al. 2019; Brecka et al. 2020) and wildlife habitats (Tremblay et al. 2018; Cadieux et al. 2019, 2020).

Although alterations of northern forest ecosystems are projected to be significant, strong spatial heterogeneity in the severity, the nature and direction of the impacts is to be expected. Such heterogeneity would likely result from the complex interactions between climate change and natural and anthropogenic disturbances, forest productivity, seed dispersal and biophysical variables (e.g., soil conditions). Disturbance severity and types strongly vary across the temperate to the boreal forest biomes and as such, are likely to differently impact successional pathways, notably when interacting with climate change. For instance, annual burned area is projected to strongly increase in central and western Canadian boreal forests when compared to eastern boreal forests (Boulanger et al. 2014), resulting in different impacts on productivity, biomass accumulation and successional pathways (Boulanger et al. 2016; Cadieux et al. 2019). Moderate and potentially selective disturbances (e.g., host-selective insect outbreaks) occurring at the temperate-boreal ecotone could favor transitions to temperate forests while climate-induced increases in severe stand-replacing disturbances would promote either transient or permanent transitions to pioneer states within the boreal forest (Payette and Delwaite 2003, Johnstone et al. 2016; Brice et al. 2020). Important modifications in species composition are to be expected at the transition zones between boreal and temperate biomes where several tree species are currently reaching either their southward or northward thermal tolerance (Leithead et al. 2010; Brice et al. 2019). On one hand, one could expected that projected increases in temperatures might favored the northward migration of e.g., thermophilous hardwood species at the expanse of boreal conifers, leading to a slow but steady northward progression of the northern hardwood forest region either through increased recruitment or better growth conditions (Duveneck et al. 2014). On the other hand, more suitable growth conditions as projected in the short-term or under mild climate forcing (D’Orangeville et al. 2018) could favor boreal species productivity in northern locations leading to faster and larger biomass accumulation as well as to closed-crown forest encroachment in sites where productivity is currently climate-limited. Despite suitable recruitment and growth conditions, propagule availability might restrain the expansion of warmed adapted species, making virtually impossible for these species to keep pace with the northward displacement of isotherms which would introduce a migration lag in the southernmost part of the boreal forest (Sittaro et al. 2017; Taylor et al. 2017). Furthermore, unsuitable, acidic soil conditions within the boreal forest could also restrain the northward expansion of hardwood species (Brown and Velland 2014) although recent analyses showed that soil characteristics represent a rather minor impediment to such transitions (Brice et al. 2020). Yet, it is still unknown how these agents of change will ultimately cumulate and interact at multiple spatial and temporal scales to alter forest landscapes over areas experiencing distinct current and future environmental conditions.

In commercial forests, anthropogenic impacts such as harvest are likely to cumulate with climate change to further modify forest ecosystems (Boulanger et al. 2016; Cadieux 2019). Harvest-induced increases in disturbance rates might modify the system’s inertia and accelerate climate-induced changes by removing resident communities and providing resources for warmed-adapted establishment and growth (Steenberg et al. 2013; Brice et al. 2020). Indeed, it was shown that alike canopy gaps (Leithead et al. 2010), moderate harvest-induced disturbance rates under warmer temperatures could favor co-occurring thermophilous pioneer but also shade-tolerant species, and hasten their northward migration within the boreal forest (Steenberg et al. 2013; Brice et al. 2019, 2020; Mina et al. 2020). When cumulated with increasing natural disturbance rates, harvest could bring total disturbance rates outside their range of natural variability (Bergeron et al. 2006; Cyr et al. 2009), resulting in regeneration failures and long-term changes in successional pathways (Splawinski et al. 2019). By favoring structural and compositional diversities, ecosystem-based forest management was suggested to promote forest resilience under increasing climate warming (Duveneck et al. 2014, 2015a; Boulanger et al. 2019). Still, this business-as-usual strategy that is now extensively applied throughout northern forests might not be efficient in order to keep or restore functional diversity under climate change (Boulanger et al. 2019; Mina et al. 2020). It has to be assessed how such a strategy might interact with climate-induced changes and how it could affect forest inertia within different forest ecosystems that will experience contrasting disturbance- and climate regime changes. In this context, quantifying the effects as well as the importance of harvest-induced impacts relative to those generated by climate change over different forest regions should offer new insights about potential adaptation avenues (Messier et al. 2019).

Evaluating the spatial heterogeneity of future climate- and harvest-induced changes in forest ecosystems is paramount in order to understand how specific drivers of forest changes are projected to proceed. Furthermore, estimating the rate and nature of the shifts as well as their overall impacts on harvest would help identify efficient and specific adaptation measures, notably to ensure the continuous provision of forest ecosystem services. Harvest and anthropogenic climate interactions on forest landscapes might be realistically projected using spatially-explicit process-based model simulations. Forest landscape models (FLM) were proven successful in order to project future changes in forest composition and structure according to different climate and harvest scenarios (e.g., Scheller and Mladenoff 2008; Steenberg et al. 2013; Duveneck et al. 2015a, 2015b, Boulanger et al. 2019). Yet, FLM simulations are frequently conducted over territories spanning less than 10M ha (with few notable exceptions, see Wang et al. 2014) which impeded the consideration of the heterogeneity in climate- and harvest-induced changes over a wide range of environmental conditions.

In this study, we modeled the impacts of harvest and increased anthropogenic climate forcing on the entire commercial forest (54.3M ha) of Quebec, Canada by using the LANDIS-II FLM. Simulations were conducted in four forest regions (northern hardwood, mixedwood, boreal east and boreal west) in eastern Canada and covered one of the largest area ever simulated with a FLM. More specifically, our objectives were to i) assess how and how much forest landscapes in each of these forest regions will be specifically altered under different climate scenarios. Furthermore, ii) we explored how harvest in the context of climate change, might differently affect forest inertia and the nature of changes projected within those forest regions. We hypothesized that i) climate change will severely altered forest landscapes with large changes driven by fire in the boreal forest regions while changes in productivity will favored hardwoods over coniferous species especially in the mixedwood region. Also ii) harvest will accelerate climate induced-changes by favoring species turnover.

## 2. Methods

### 2.1 Study regions

We projected forest landscapes over the entire commercial forest in the province of Québec (54.3 M ha, Figure 1). The area is bordered on the southeast by the northern extent of the Appalachian Mountains. In this area, valleys covered by glacial till and humo-ferric podzols are intermingled with broadly rolling mosaic of upland plateaus sitting between 200 and 800 m above sea level. Most of the area located north of the St. Lawrence River belongs to the Canadian Shield. Landform and soils in this area are dominated by uplands and wetlands, where Precambrian granitic bedrock outcrops alternate with ridges encompassing coarse texture hummocky deposits of glacial origin. Climate is more typical of the temperate continental region in the southernmost part of the study area, whereas the northernmost part is typical of boreal regions. The study area covers wide latitudinal and longitudinal temperature gradients (mean annual temperature south MAT = 6.6°C; north MAT = −3.1°C and total annual precipitations west TAP = 600 mm; east TAP = 1200 mm, Robitaille and Saucier [1998]). A wide variety of forest ecosystems occur within the area and as such, the study area was divided in four forest regions based on the Quebec’s bioclimatic subdomains, i.e., the northern hardwoods, the mixedwoods as well as the boreal east and boreal west (Robitaille and Saucier 1998). Species-rich northern hardwood forests dominated by deciduous mesophytic species are located in the southern part of the study area. Forest gradually transitions to mixed in the mixedwood forest region and finally to conifer-dominated forests with increasing latitude in both boreal forest regions. Recurrent spruce budworm (SBW) outbreaks are the most important natural disturbance in the mixedwood forest region (Boulanger et al. 2012), and small windthrows and single-tree mortality drive natural forest succession in the northern hardwood forest region. Wildfires are most prevalent within the boreal portion of the study area with fire return intervals decreasing from *ca* 400 years in the east to *ca* 100 years in the drier, western part (Boulanger et al. 2014). Timber harvest occurs at various rates over the entire study area, with cutblock size and proportion of biomass harvested typically increasing with latitude. Single-tree and small-patch harvest are most prevalent in the northern hardwoods, whereas clearcuts reaching more than 100 ha are more common in the boreal regions. These prescriptions follows ecosystem-based forest management guidelines and aim at emulate natural disturbances in each of these regions.

**Figure 1.**
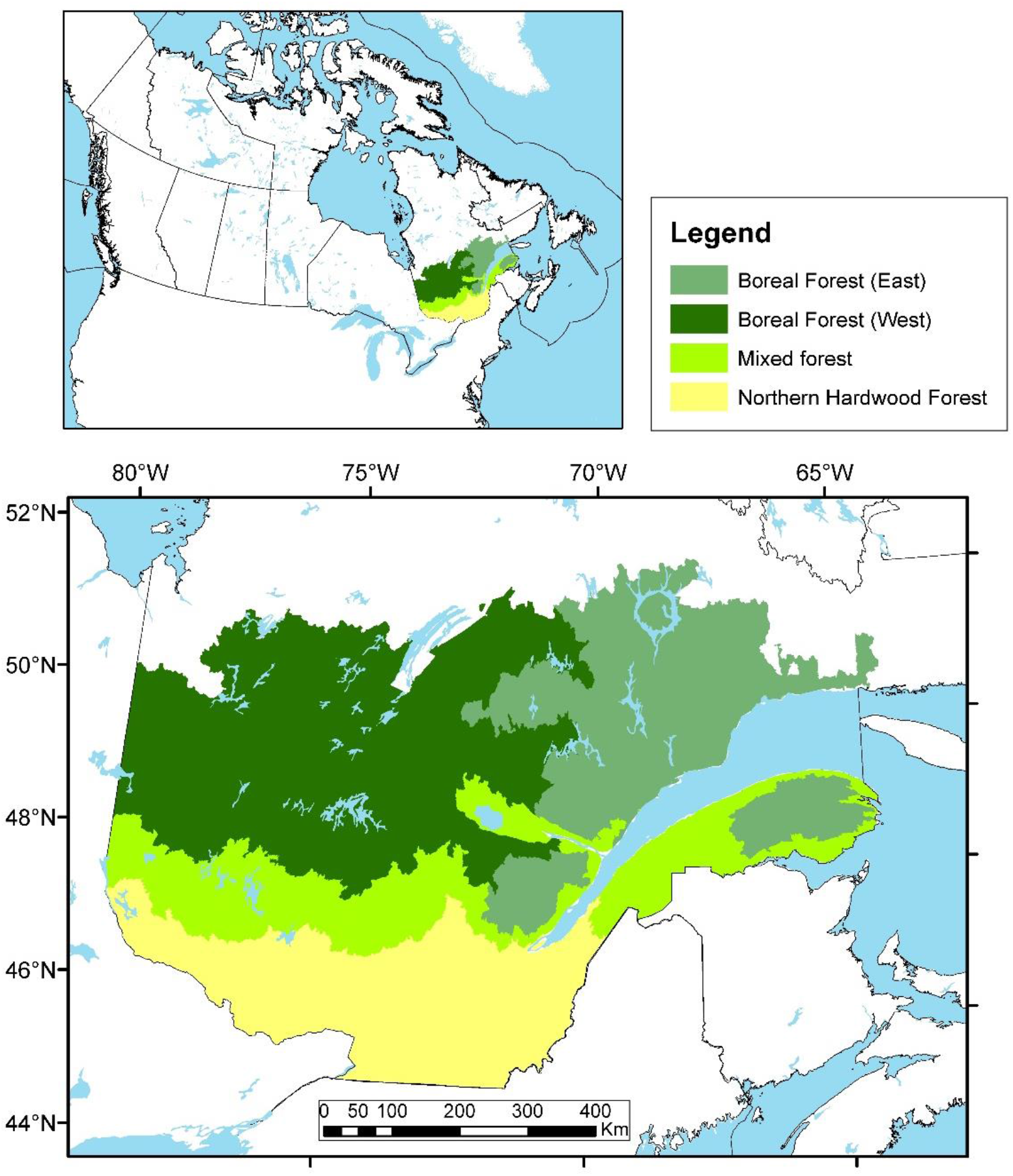
Location of the study area as well as the four forest regions considered for analyses.

### 2.2 Climate data

We produced monthly time series for baseline climate by interpolating data from climate station records (McKenney et al. 2013). We obtained future climate projections from the Canadian Earth System Model version 2 (CanESM2) for each of two different radiative forcing scenarios, i.e. RCP 4.5 and RCP 8.5. Radiative forcing is assumed to stabilize at 4.5 W·m^−2^ after 2100 in the RCP 4.5 scenario, without an “overshoot” pathway. In the RCP 8.5 scenario, the forcing reaches 8.5 W·m^−2^ in 2100 and keeps on increasing afterwards. Under these scenarios, the CanESM2 projects mean annual temperature increases of ca 3.9°C (RCP 4.5) to ca 8.5°C (RCP 8.5) throughout the study area by 2100 (compared to circa 2000), while average precipitation is projected to increase by 7% (RCP 8.5) to 10% (RCP 4.5). We bias-corrected data from CanESM2 for the 1900 - 2100 period by expressing temperatures as differences and precipitations as ratios relative to the CanESM2 monthly means for the 1961-1990 period.

### 2.3 LANDIS-II forest landscape simulation model

LANDIS-II is a spatially-explicit raster-based forest landscape model that simulates disturbances, seed dispersal, and forest succession (Scheller et al. 2007). Species are defined using unique life-history attributes and are represented in each grid cell as 10-year age-cohorts. Forest composition and structure in each cell were initialized using provincial ecoforestry maps and cohort data from provincial permanent and temporary forest inventory plots (FIP). These maps were rasterized at a 250-m (6.25 ha) resolution. Each of these cells was then assigned to a spatial unit (i.e., “landtype”) with homogeneous soil and climate conditions. Grid cells with more than 50% of their area covered with non-forest cover types were classified as inactive.

### 2.4 Forest succession and species growth potential

We used the LANDIS-II Biomass Succession extension v 3.2 (Scheller and Mladenoff 2004) to simulate forest succession in each 250-m cell. This extension simulates modifications in cohort aboveground biomass (AGB) over time by taking into consideration tree species’ cohort age, life-history traits, and species-specific landtype responses (Suppl. Mat. S1). We used species’ life-history traits information collected from various sources including several previous LANDIS-II publications conducted for North American forest landscapes. Species were classified as either thermophilous or boreal according to their thermal preference in growing degree-days (Suppl. Mat. S1). We also classified species according to which successional stage, e.g., either pioneer or mid-/late-successional, they are mostly associated consistent with their shade tolerance and longevity.

We parameterized and calibrated three sets of dynamics inputs sensitive to soil and climate conditions, i.e., i) species establishment probabilities (*SEP*), ii) maximum possible aboveground net primary productivity (*maxANPP*), and iii) maximum aboveground biomass (*maxAGB*). Parameterization was conducted using the individual tree-based, forest patch model PICUS version 1.5 (Lexer and Honninger 2001; Taylor et al. 2017). PICUS simulates the dynamics of individual trees on 10×10 m patches across forest stand areas and accounts for spatially-explicit interactions among patches via a 3D light module. PICUS simulates the effects of climate and soil properties on tree population dynamics (Lexer and Honninger 2001). Using individual tree species parameters, we ran PICUS simulations for 17 tree species occurring in the study regions (Suppl. Mat. S1). A complete description of the model and how it was parameterized and validated can be found in Taylor et al. (2017). In order to determine the three dynamic input parameters for Biomass Succession extension, we simulated mono-specific 1-ha stands with PICUS for each of the 17 tree species. A factorial simulation design was used to simulate all mono-specific stands for tree species and landtype, under climate conditions for specific periods (2000-2010, 2011-2040, 2041-2070, 2071-2200) and forcing scenarios (baseline, RCP 4.5, RCP 8.5). Simulations were run for 300 years, starting from bare-ground and used the landtype-specific soil and the period- and climate scenario specific climate data. Values for *SEP*, *maxANPP* and *maxAGB* were then derived from these simulations (see Boulanger et al. 2016). Previous analyses (Boulanger et al. 2016, 2017, 2019) have shown good agreement between successional pathways predicted by LANDIS with those reported in the literature. Readers can refer to Supplementary Material S2 for *maxAGB* maps.

### 2.5 Forest harvest

The Biomass Harvest extension (v3.0; Gustafson et al. 2000) was used to simulate forest harvest. Relevant information regarding harvest parameters such as mean harvested patch size and total harvested area, was summarized by forest management units (FMU). Harvest was set to vary according to potential vegetation as defined by the Quebec’s Hierarchical System for Territorial Ecological Classification (Bergeron et al. 1992). This system classifies forest stands according to their potential natural vegetation type, which is a function of climatological and geomorphological constraints on vegetation growth and succession. We simulated ecosystem-based forest management (EBFM), a harvest scenario that should closely mimic the historical disturbance regimes (wildfires, spruce budworm outbreaks, single-tree mortality, small gap openings) occurring in the study area. Rotation length time and biomass removal levels were fixed according to current harvest regulations and expert advices as in Boulanger et al. (2019). Harvest rates were held constant throughout the simulations unless not enough stands qualified for harvest. In this latter case, harvest proceeded until there were no more stands available.

### 2.6 Natural disturbances

Fire, and spruce budworm (SBW, *Choristoneura fumiferana* [Clem.]) outbreaks, were considered as natural disturbances in the LANDIS-II simulations. Both disturbances historically had major impacts on Quebec’ forest landscapes (e.g., Bouchard et al. 2006). SBW outbreaks are mostly prevalent within the mixedwood region whereas fires are more important within the boreal regions. The LANDIS-II Base Fire v3.0 extension was used to simulate stochastic fire events dependent upon fire ignition, initiation and spread. Fire regime data (annual area burned, fire occurrence, and mean fire size) were summarized into “fire regions” corresponding to the intersection of the study area and the Canadian Homogeneous Fire Regime zones of Boulanger et al. (2014). Baseline and future fire regime parameters within each fire region were calibrated according to models developed by Boulanger et al. (2014) and further updated for different RCP scenarios (Gauthier et al. 2015). Annual area burned (AAB) will remain minimal within northern hardwoods, regardless of climate scenarios whereas it will reach maximum values > 2% after 2080 in boreal west under RCP 8.5 (Supplementary Material S3).

The Biological Disturbance Agent (BDA) v3.0 extension (Sturtevant et al. 2004) modified to account for specific SBW parameters was used to simulate SBW outbreaks. From the most to least vulnerable, host tree species for SBW included balsam fir (*Abies balsamea*), and white (*Picea glauca*), red (*P. rubens*) and black (*P. mariana*) spruces (Hennigar et al. 2008). Outbreaks are simulated as probabilistic events at the cell level with probabilities being a function of the site and neighborhood resource dominance (e.g., host species occurrence within a 1-km radius) as well as regional outbreak status. Outbreak-related tree mortality is contingent on these probabilities as well as on host species- and age-specific susceptibility. SBW outbreak parameters were calibrated and validated using various studies conducted within the boreal and mixedwood forests (Hennigar et al. 2008). Regional outbreaks were calibrated at the highest severity level possible were set to last one-time step (10 years) and to recur every 40 years in accordance with historical regional cycles (Boulanger et al. 2012).

### 2.7 Simulation design

As the study area is one of the largest ever simulated with LANDIS-II, we had to split it into five sub-areas that were simulated separately for computational reasons. In order to consider some potential edge effect related to these zones pertaining to e.g., seed dispersal or fire spread from nearby regions, we simulated each sub-area with an additional 50-km buffer overlapping the adjacent sub-areas. Simulations were run according to a factorial design, i.e., under the three climate projections, (corresponding to baseline, and the RCP 4.5 and RCP 8.5 radiative forcing scenarios) and two harvest scenarios. The two harvest scenarios were no harvest and a baseline, EBFM harvest where current parameters described above were applied. Five replicates were run for 130 years for each harvest and climate change scenario combination, starting in the year 2020, and using 10-year time steps. Except for scenarios involving the baseline climate, we used the projected fire regime parameters projected for the 2011-2040 periods for the 2020-2040 simulated years. Also, fire regime parameters were allowed to change in 2041-2050 and 2071-2080 according to the average climate corresponding to each forcing scenario. Fire regime parameters for 2071-2080 were held constant up to 2150. Dynamic growth and establishment parameters (*SEP, maxANPP* and *maxAGB*), were allowed to change according to each climate scenario following the same schedule used for the fire regime parameters.

### 2.8 Analyses

We tested for the impacts of climate change and current harvest practices on forest landscape by calculating dissimilarities between projected forest landscapes and those under baseline climate. Analyses were performed by calculating Bray-Curtis dissimilarity (d_BC_) using species-specific AGB. First, species-specific AGB was averaged at the landtype-level at each timestep among replicates. Dissimilarities were then calculated relative to projections at time *t* as well as according to the harvest scenario under each climate forcing scenario separately. Landtype-level dissimilarities were then area-weight averaged at the forest region level.

Dissimilarities between projected and baseline forest landscapes can arise from changes in species-specific abundance (AGB), from changes in species composition *per se*, or from both (Basega 2013). Separating these components helps to identify the empirical patterns underlying changes in species composition (Basega 2013). We therefore decomposed all d_BC_ values into their two additive components i.e., the fraction linked to balanced variation in abundance (d_BC-bal_) and the one linked to abundance gradients (d_BC-gra_) (Basega 2013). d_BC-bal_ informs about change in species composition or species turnover whereas d_BC-gra_ is related to change in species abundance. We then assessed trends in these two components according to climate and harvest scenarios in each of the four forest regions. All d_BC_ components were calculated using the *betapart* v1.5.2 package (Basega et al. 2020) in R 3.6.2 (R Core Team 2019).

## Results

### Climate change impacts across forest regions

Generally speaking, dissimilarities were highest for boreal west and appeared sooner in the simulations while being lowest for the northern hardwood throughout the simulated period, regardless of the climate scenarios (Fig. 2). That being said, dissimilarities increased over time and with increasing anthropogenic climate forcing in all regions (Fig. 2, see also Suppl. Mat S4). Dissimilarities under RCP 4.5 and RCP 8.5 strongly diverged in all regions in 2080-2100, showing a sharp increase under RCP 8.5 (Fig. 2).

**Figure 2.**
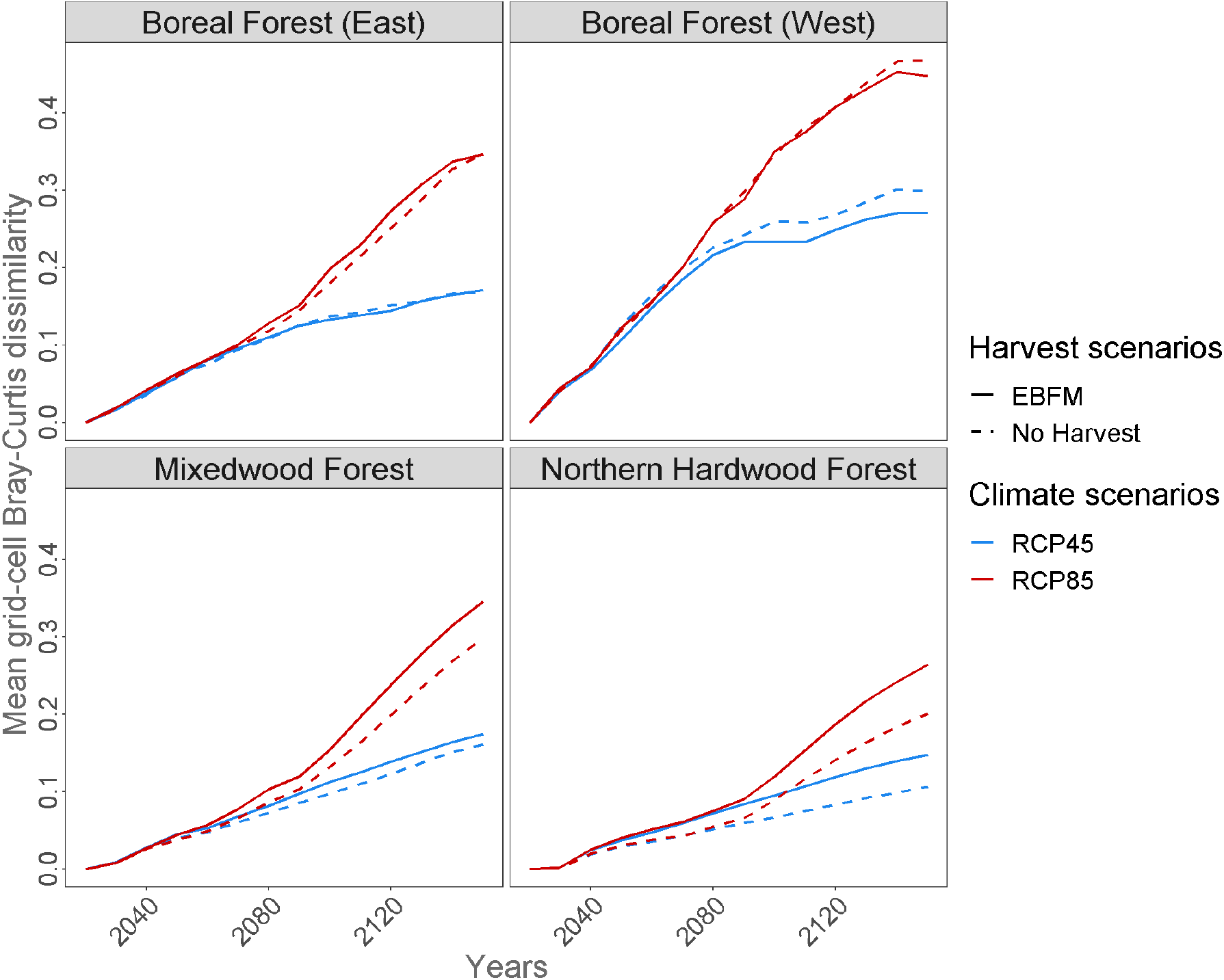
Temporal trends in Bray-Curtis dissimilarities (d_BC_) under RCP 4.5 and RCP 8.5 and the two harvest scenarios (EBFM = Ecosystem-based forest management; No harvest). Dissimilarities were calculated against the respective harvest scenario under baseline climate.

With very few exceptions, dissimilarities with baseline climate forest landscapes were mostly linked to gradients in abundance (d_BC-gra_ > 50%) before 2100 rather than changes in species composition (d_BC-bal_) *per se*, regardless of forest regions and simulation scenarios (Figure 3). However, there were important differences in d_BC-bal_ trends between regions and scenarios. d_BC-bal_ tended to be higher in the mixedwood and northern hardwood forest regions, especially under RCP 4.5, while being lowest in the boreal west where it never exceeded 30% (Figure 3). In 2150 under RCP 4.5, more than 50% of the dissimilarities were related to d_BC-bal_ in the northern hardwood and mixedwood forest regions. Under RCP 8.5, d_BC-bal_ never exceeded 50% in 2150 in any regions and was highest in the mixedwood and boreal east (Figure 3). d_BC-bal_ tended to increase with time in all scenarios except under RCP 8.5 in both mixedwood and northern hardwood regions. In these regions, d_BC-bal_ first increased up to 2080-2100. From this point, d_BC-bal_ sharply dropped under RCP 8.5 while it continued to increase under RCP 4.5 (Figure 3).

**Figure 3.**
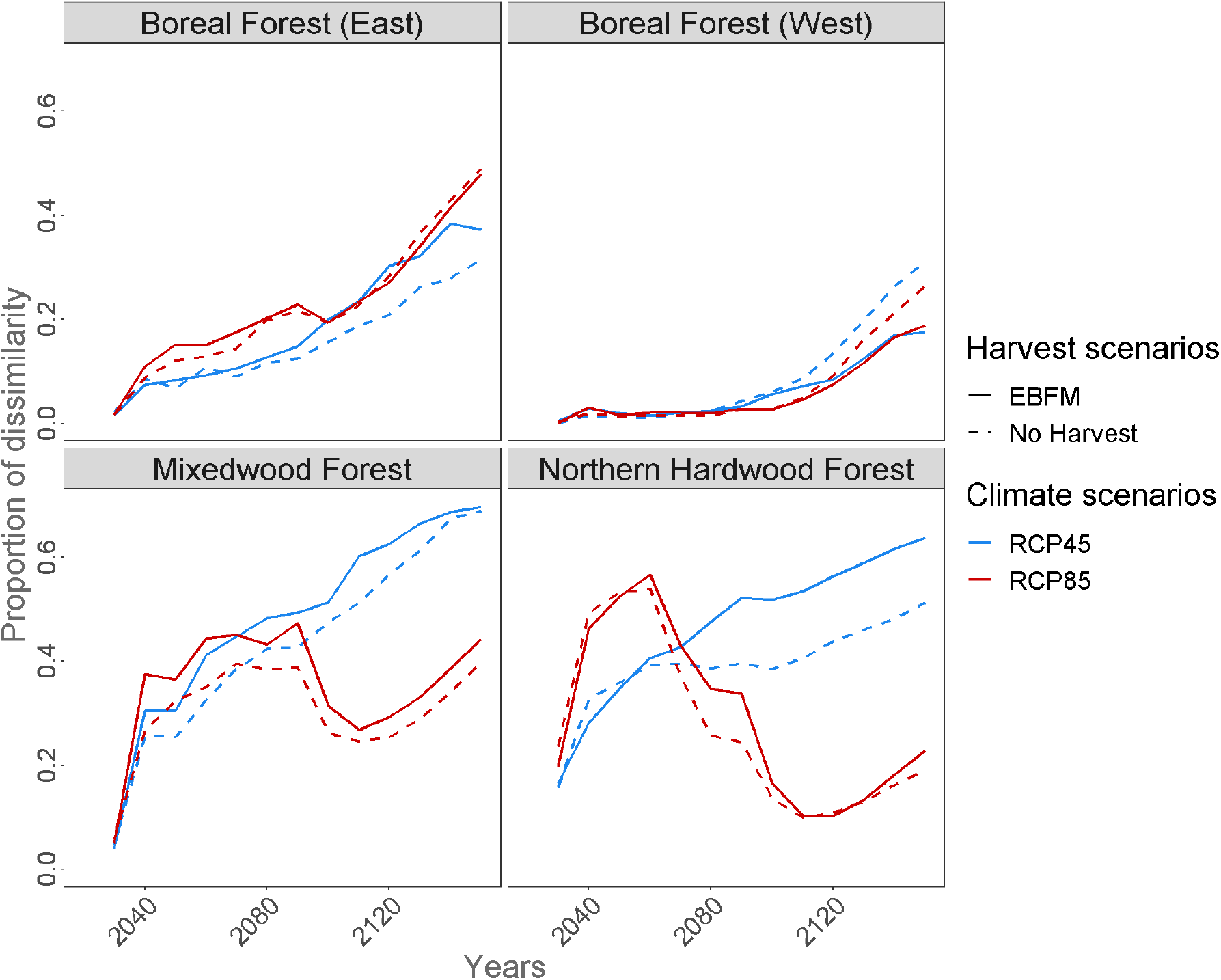
Temporal trends in the proportion of Bray-Curtis dissimilarities shown in (figure 2 linked to balanced variation in species abundance (d_BC-bal_, Baselga 2013). Recall that dBC-bal is the inverse of the variation in Bray-Curtis dissimilarity linked to abundance gradient, i.e, d_BC-gra_ = 1-d_BC-bal_ under RCP 4.5 and RCP 8.5 and the two harvest scenarios. Results are shown for the two climate and two harvest scenarios separately.

Dissimilarities linked to gradient in abundance can be associated with the strong climate-induced decline in total AGB throughout the study under RCP 8.5, most notably after 2100 (Fig. 4). In boreal west, these declines would occur much sooner, i.e., as soon as 2050 under milder (RCP 4.5) anthropogenic climate forcing. Under RCP 8.5, the total decrease in AGB relative to baseline climate was highest in this latter region (−54%) whereas AGB decline in other regions would reach between 35 and 40% by 2150 (Fig. 4). AGB decreases under RCP 4.5 would be much lower in any region. Under this scenario, total AGB in mixedwood and northern hardwood forest regions would decrease by 10 to 15% relative to baseline climate while differences in AGB between RCP 4.5 and the baseline climate would be virtually nil (Fig. 4).

**Figure 4.**
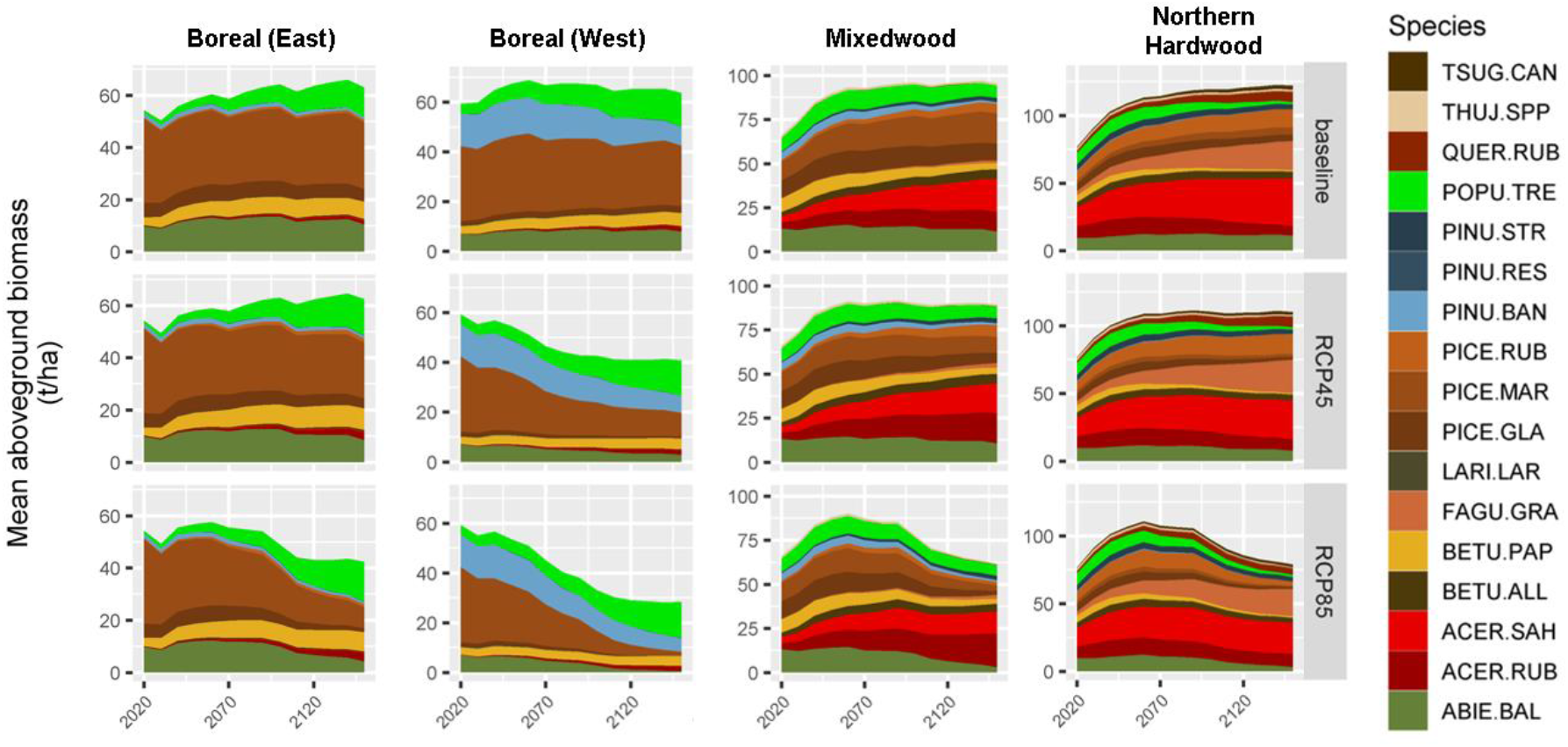
Stacked trends in species AGB for each of the four forest regions simulated under either baseline, RCP 4.5 or RCP 8.5 climate scenarios. See Suppl. Mat. S1 for species abbreviations. Only simulations considering baseline harvest are considered here.

In all forest regions, decrease in total AGB is mostly associated with important declines in boreal conifer AGB (Fig. 4, Suppl. Mat S5). These declines were more important with increasing anthropogenic climate forcing and with decreasing latitude. AGB declines were particularly important for white and black spruces regardless of the climate scenarios (Fig. 4). More notably, declines in black spruce in boreal west, which is by far the most common species under current conditions in this region, would be dramatic as it would almost completely disappear by 2150 under RCP 8.5 (Suppl. Mat S5). Balsam fir and red spruce would also decline throughout the study area but mostly under RCP 8.5. Throughout the study area, climate-induced decreases in boreal conifer AGB would not be compensated by an increase in thermophilous species biomass which would rather maintain stable biomass relative to baseline climate (Fig. 4). Notable exception includes red maple AGB that would mostly increase under RCP 8.5 especially in the boreal east and mixedwood forest regions (Fig. 4).

As such, higher d_BC-bal_ in southernmost forest regions can be interpreted as gradual species turnovers from boreal to thermophilous species (Fig. 5). Increases in thermophilous proportions were highest in the mixedwood forest region (8 – 25% depending on scenarios with increased forcing) and were likely due to an increase in the proportions of red and sugar maples as well as American beech (Fig. 4). Overall increase in thermophilous proportions were relatively small (< 10%) in both boreal forest regions compared with baseline climate (Fig. 5). In these regions, changes in species composition were mostly related to transition from mid-/late successional species to higher proportions of pioneer species (Fig. 6). Pioneer species proportions strongly increased relative to baseline climate in boreal (+10-50%) but also in mixedwood (+10-25%) forest regions while this increase was much smaller in the northern hardwood region (Fig. 6). Increases were particularly important under RCP 8.5, notably after 2100 and in the boreal west region. Increases in pioneer proportions were mostly resulting from increased in trembling aspen (boreal regions) and red maple (mixedwood) AGB (Fig. 4).

**Figure 5.**
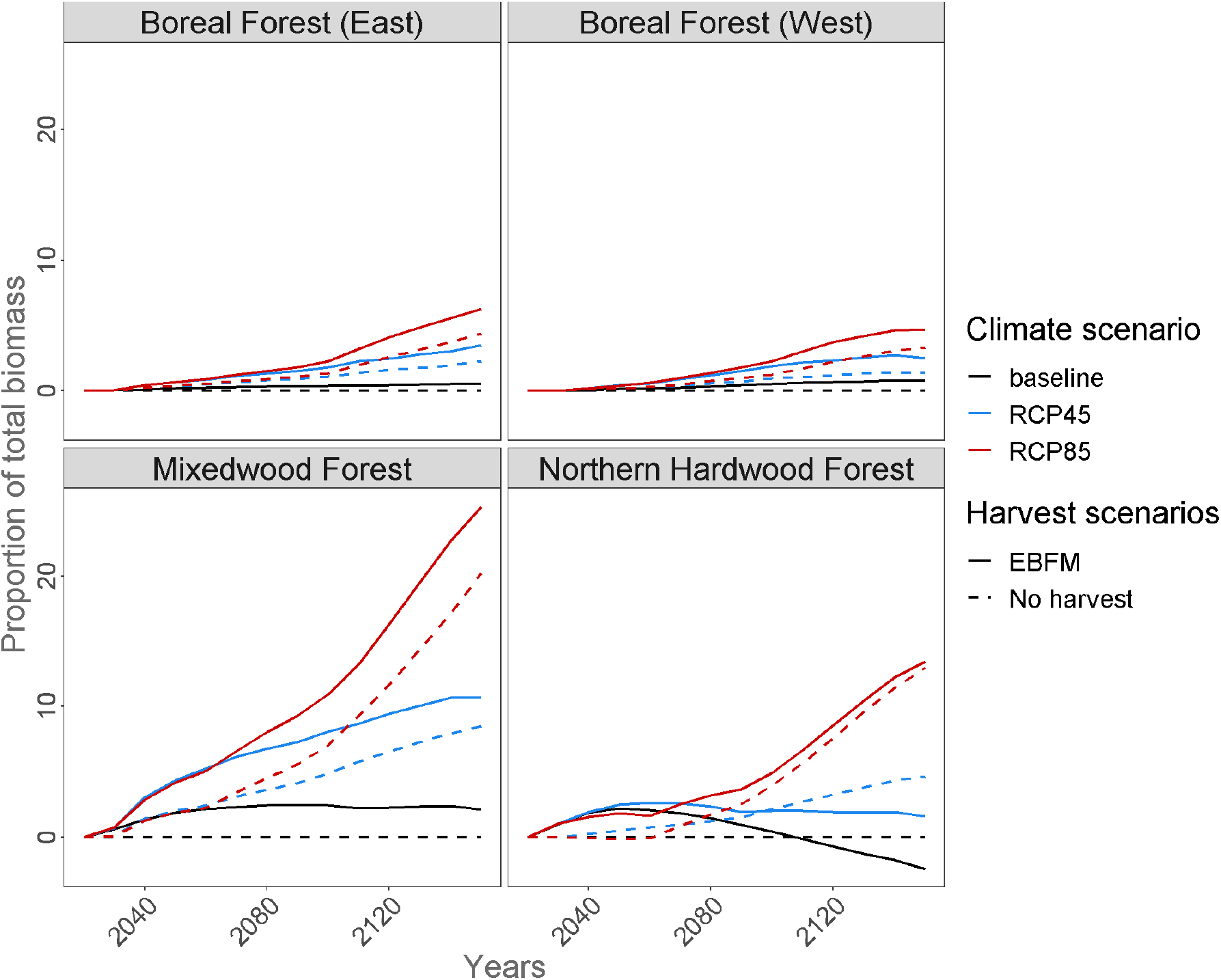
Temporal trends in proportions of thermophilous species (see table 1 for definition) under the three climate and two harvest scenarios. Results are expressed as differences with simulations conducted under baseline climate and no harvest at time *t*.

**Figure 6.**
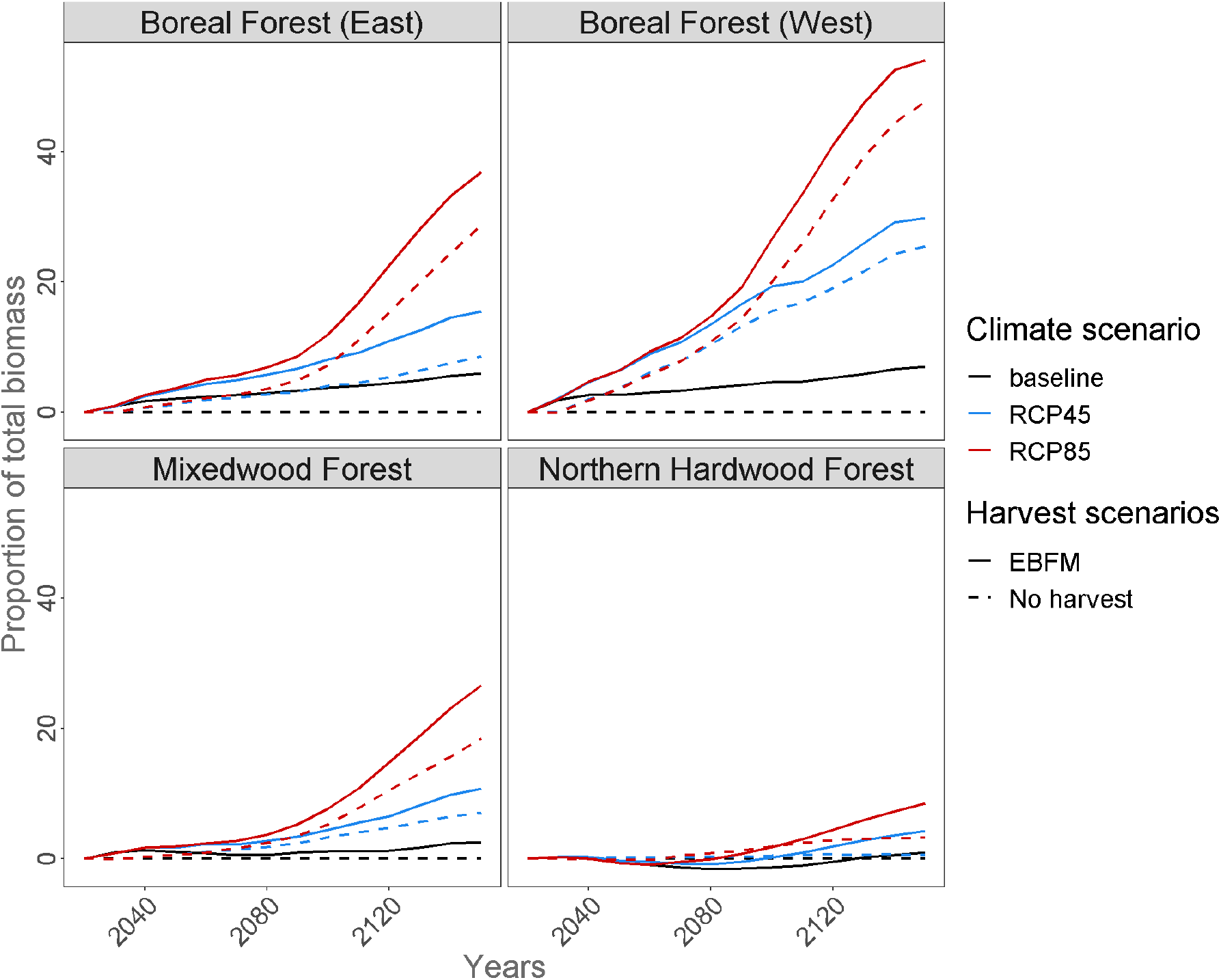
Same as figure 5 but for pioneer species proportions.

### Harvest impacts in the climate change context

Harvest was projected to generate more dissimilarities (higher d_BC_) between projected forest landscape and those under baseline climate than when it is not included in the simulations, for both the mixedwood and northern hardwood forest regions, regardless of the climate scenario (Figure 2, see also Suppl. Mat S2). Although overall d_BC_ were low, harvest had the most important impacts on dissimilarities in northern hardwoods when compared with scenarios without harvest (Figure 2). As opposed, in both boreal regions, virtually no difference in dissimilarities with baseline climate were projected between scenarios with or without harvest with the exception of boreal west under RCP 4.5. Moreover, harvest had little impacts on how climate induced dissimilarities with baseline forest landscapes. Indeed, difference in d_BC-bal_ values were rather small (Figure 3) with d_BC-bal_ tending to be slightly higher under harvest than under no harvest.

Harvest promoted and accelerated the proportion of thermophilous species AGB within the mixedwood forest region regardless of climate scenario, but most notably under RCP 8.5 (Fig. 5). In this region, thermophilous species proportions were at least 5 points of percentage higher with harvest than no harvest under RCP 8.5. That being said, thermophilous species also expanded with harvest when compared to simulation without harvest, but at lower rates, under no or milder anthropogenic climate forcing in this forest region. Harvest impacts in this regard were relatively small in other forest regions (+1-2 points of percentage). As expected, harvest also promoted pioneer species AGB virtually everywhere (Fig. 6). Impacts were slightly more important under RCP 8.5 than under RCP 4.5 in the mixedwood and boreal west forest regions.

## Discussion

Our results showed that climate change impacts will be striking in eastern Canada forest regions. Simulated changes were most important within the boreal forest regions, where they mainly translated into very sharp decreases in AGB. Important increase in wildfire activity which could reach >2 % of area burned per year by 2100 under RCP 8.5 (Suppl. Mat. S3) could explain these declines, most notably in boreal west. Increased annual area burned will result in extensive conversion of old-growth boreal-dominated stands into regenerating and young stands, which comprise much lower AGB (Boulanger et al. 2016). Very short fire intervals (< 30 years) under RCP 8.5 in boreal west could also extensively preclude the regeneration of non-pyrophilous and late-successional species such as white spruce and balsam fir, further contributing to this decline. Most importantly, such an increase in wildfires could also impede black spruce regeneration by preventing individuals to reach sexual maturity. Overall, such declines might cause large areas in eastern Canada’s boreal forests to experience “regeneration failures” which would further decrease the long-term ability of the boreal forest to accumulate AGB (Gauthier et al. 2015; Splawinski et al. 2019). This result is particularly dramatic as it could lead to the widespread extirpation (Suppl. Mat. S5) of black spruce, the Canada’s boreal flagship species, in boreal west. As a corollary, increased disturbance rates under more important climate forcing will favour pioneer species, most notably trembling aspen, which were projected to mostly expand in boreal regions. Aspen was shown to increase with more frequent anthropogenic disturbances, including fires, within the last 200 years in eastern Canada’s boreal forest (Danneyrolles et al. 2016) and is also projected to strongly increase with climate-induced changes throughout the boreal forest (Boulanger et al. 2016, 2017; Brecka et al. 2020).

Increased natural disturbance rates within the boreal forest regions likely cumulated with climate-induced growth constraints to most boreal species to further reduce total AGB. Strong warming, especially under RCP 8.5, will gradually reduce the potential productivity of several boreal species throughout eastern Canada, except in the northernmost and most elevated lands (Suppl. Mat S2), as species will experience conditions further from their optimal climate tolerance. Similar declines in productivity have also been projected for boreal species within most of their range under aggressive climate forcing (Girardin et al. 2016; D’Orangeville et al. 2019), primarily as a result of increased metabolic respiration following more frequent and severe drought conditions (Girardin et al. 2016). Lack of propagules of warm-adapted species within most of the boreal range impeded potential AGB recovery from boreal conifer decline. In fact, thermophilous species migration towards more northern, suitable conditions will be strongly outpaced by the northward shift of the isotherms even under moderate forcing (Brice et al. 2019; Prasad et al. 2020). Considering their limited dispersal abilities, we projected that thermophilous species would colonize sites only few tens of km north of their actual northern range (Suppl. Mat S5), although suitable conditions would expand much further north under increased climate forcing (Suppl. Mat S2). Additionally, some thermophilous species (e.g., sugar maple) might face important growth constraints, presumably drought-induced (Taylor et al. 2017) even north of their current range under aggressive climate forcing (Suppl. Mat S2), compromising their potential expansion within the boreal forest. Notwithstanding the increased disturbance rates, thermophilous species are therefore less likely to extensively compensate AGB losses by boreal species by introducing a significant “migration lag” within the boreal forest (Taylor et al. 2017; Prasad et al. 2020).

In the more southern mixedwood and northern hardwood regions, changes will be less important than within boreal regions and will further result from alteration in species composition. Thermophilous species proportions within those two forest regions will strongly increase at the expanse boreal species. Climate-induced changes in growth directly altered the competitive abilities of boreal species, making these more likely to be outperformed by co-occurring thermophilous species (Fisichelli et al. 2014; Reich et al. 2015; Boulanger et al. 2016). As opposed to boreal forest regions, this co-occurrence allows thermophilous species to partially compensate the decline in boreal species AGB in the mixedwood and northern hardwood. Notably, reduced climate-induced recruitment for balsam fir following SBW outbreaks could have allowed for co-occurring hardwood species turnover, leading to permanent thermophilization of the ecotone (Brice et al. 2019, 2020). Combined with the boreal migration lag mentioned above, this suggests that the mixedwood forest region might strongly contract in the upcoming decades, making the ecotone between the northern hardwood and boreal forest much more abrupt.

Our results suggested that the boreal forest regions will be much less resilient than northern hardwoods to the projected changes in climate regimes. The boreal forest is known to have been particularly resilient to variations in climate and disturbance regimes during the last several millennia (Carcaillet et al. 2010). Indeed, stand-replacing disturbances (e.g., fires) that are mostly occurring within the boreal forests were hypothesized to promote resilience by favoring the recovery of resident species (Liang et al. 2018). However, projected disturbance rates in the boreal forest will likely fell outside the range of natural variability, triggering potential important ecosystem shifts (Bergeron et al. 2006). Cumulative impacts of wildfires and productivity decline within boreal west could result in alternative stable states (Stralberg et al. 2018) toward landscapes more reminiscent of open parklands or taiga (Girard et al. 2007). Therefore, this extensive disturbance- and climate-induced biome shift projected for the next century would be unprecedented since the last glaciation in this area. Similar transitions from boreal forest to open parklands or prairie ecosystems is also projected for several regions in western Canada as a result of increase drought and wildfire activity (Stralberg et al. 2018). Diversified forests were shown to be more resilient under increased anthropogenic climate forcing (Duveneck et al. 2014; Mina et al. 2020). Higher species diversity along with diversified functional traits and low projected disturbance rates will help northern hardwood forest landscapes being more resilient to climate-induced changes although species turnover will likely occur.

Our analyses showed that current harvest strategies will strongly decrease forest inertia and will interact with anthropogenic climate forcing to further modify forest landscapes notably in the southernmost forest regions. In the mixedwood forest region for instance, harvest will accelerate climate-induced changes by hastening the increase in thermophilous species AGB, promoting a more rapid biome shift toward communities more typical of the northern hardwoods than under no harvest. Increased disturbance rates are known to catalyze changes toward more adapted species (Thom et al. 2017). We showed that harvest will favour opportunistic species such as red maple, a species known as a “supergeneralist” for which historical harvest already contributed to its swift expansion throughout northeastern North America (Danneyrolles et al. 2016), notably by interacting with climate change (Brice et al. 2020). Harvest strategies likely played a role to affect forest inertia differently between forest regions. Frequent and widespread partial cutting (typically removing ~25-40% of total biomass over > 3% of the territory per year) in the mixedwood and northern hardwood, as oppose to relatively infrequent but severe clearcuts (removing 90-100% of the biomass over < 1% of the territory per year) in the boreal forests, potentially affected forest inertia more severely over a large portion of the territory, hence accelerating species turnover in these southern regions. Moderate disturbances as these harvest strategies were shown to strongly decrease forest inertia and favor species turnover by promoting hardwood species recruitment and growth at the temperate-boreal ecotone (Brice et al. 2019, 2020). As after natural disturbances that these harvest management strategies are emulating, climate-induced growth constraints on boreal species make them less likely to recolonize small gaps in areas where they co-occur with thermophilous species (Leithead et al. 2010). Furthermore, natural disturbances are projected to remain stable and/or infrequent (wildfires, windthrow) or even to decrease (SBW outbreaks) in the mixedwood and northern hardwood (Boulanger et al. 2014, 2016). This makes harvest more likely to alter forest inertia in these regions than within the two boreal forest regions, especially boreal west, where climate impacts on disturbance rates will overwhelm those generated by harvest even in the short term.

The forest sector is currently one of the most important industry in eastern Canada, representing $11.5 MM, i.e., 4.4% of Quebec’s GDP in 2006 (MFFP 2016). Alterations in biomass, age structure and species composition resulting from changes in natural disturbances and climate-induced growth constraints would likely reduce timber supply (Gauthier et al. 2015; Daniel et al. 2017; Boucher et al. 2019; Brecka et al. 2020) which would greatly affect supply value chains (Yemshanov et al. 2018). A posteriori analyses showed that strong anthropogenic climate forcing will reduce harvested biomass by as much as 40% by 2150 (Suppl. Mat. S6). Furthermore, concurrent with the overall decline in harvested biomass, strong compositional shifts towards deciduous species in the mixedwood could have serious economic implications as conifers are generally preferred by the industry. The type and quality of wood products that companies can manufacture (Brecka et al. 2020) could thus be seriously affected. In this context, increased costs, timber supply shortages and important impacts on the supply value chains are to be expected throughout the study area, with a potential collapse of timber harvest in some regions (McKenney et al. 2016, Brecka et al. 2020). Major changes to harvest practices that consider projected wildfire activities and changes in productivity (Irulappa Pillai Vijayakumar et al. 2016; Boulanger et al. 2017) are thus strongly needed to preserve the long-term sustainability of wood supply in eastern Canada. Adaptation strategies should be region-specific as we showed that climate change will affect forest landscapes differently throughout the study area. Reducing long-term harvest targets by lengthening rotation periods could ensure steady, sustainable timber supplies by maintaining a stock of timber that could buffer the effects of wildfire activity (Raulier et al. 2014; Brecka et al. 2020). Furthermore, harvest strategies fostering natural transition to thermophilous species in the mixedwood forest region could be favored (Pedlar et al. 2012;). In combination with e.g., assisted migration, these strategies would help compensate biomass loss in regions where boreal species growth will be strongly constrained. Eastern Canada forest ecosystems might cross important tipping points leading to significant changes under strong anthropogenic climate forcing. We showed that climate-induced impacts were much more important and swifter under RCP 8.5, with great divergence with baseline-climate forest landscapes occurring after 2080 throughout the study area. We showed that significant forest landscape alterations, notably total AGB declines, might be prevented under RCP 4.5 for most of eastern Canada (except boreal west), presumably causing minimal impacts on ecosystem services. This should call for strong mitigation measures in order to maintain anthropogenic climate forcing to lower values than those expected by the end of the century under business-as-usual global strategies (Hausfather and Peter 2020).

## Acknowledgements

We want to acknowledge Jean-Daniel Sylvain who provided the soil data. We also want to thank Anthony Taylor and David Price who provided PICUS parameters and climate data respectively. Dominic Cyr helped parameterize LANDIS-II. This research was funded by Natural Resources Canada.

---

Supplementary Material S1 – LANDIS-II and PICUS parameters

**Table S1.** LANDIS-II input data for tree species simulated within the study area.

| Species | Species code | Longevity | Age at maturity | Shade tolerance† | Effective seed dispersal (m)‡ | Maximum seed dispersal (m) | Vegetative regeneration | Post-fire regeneration | Growth curve shape parameter | Mortality curve shape parameter | Thermal preference* | Successional stage** |
| --- | --- | --- | --- | --- | --- | --- | --- | --- | --- | --- | --- | --- |
| *Abies balsamea* | ABIE.BAL | 150 | 30 | 5 | 25 | 160 | No | Seeding | 0 | 25 | boreal | M/L- succ. |
| *Acer rubrum* | ACER.RUB | 150 | 10 | 3 | 100 | 200 | Yes | Resprout | 0 | 25 | Therm. | Pioneer |
| *Acer saccharum* | ACER.SAH | 300 | 40 | 5 | 100 | 200 | Yes | Resprout | 1 | 15 | Therm. | M/L- succ. |
| *Betula alleghaniensis* | BETU.ALL | 300 | 40 | 3 | 100 | 400 | Yes | Resprout | 1 | 15 | Therm. | M/L- succ. |
| *Betula papyrifera* | BETU.PAP | 150 | 20 | 2 | 200 | 5000 | Yes | Resprout | 0 | 25 | boreal | Pioneer |
| *Fagus grandifolia* | FAGU.GRA | 250 | 40 | 5 | 30 | 3000 | Yes | Seeding | 1 | 15 | Therm. | M/L- succ. |
| *Larix laricina* | LARI.LAR | 150 | 40 | 1 | 50 | 200 | No | Seeding | 0 | 25 | boreal | Pioneer |
| *Picea glauca* | PICE.GLA | 200 | 30 | 3 | 100 | 303 | No | Seeding | 1 | 15 | boreal | M/L- succ. |
| *Picea mariana* | PICE.MAR | 200 | 30 | 4 | 80 | 200 | No | Serotiny | 1 | 15 | boreal | M/L- succ. |
| *Picea rubens* | PICE.RUB | 300 | 30 | 4 | 100 | 303 | No | Seeding | 1 | 15 | boreal | M/L- succ. |
| *Pinus banksiana* | PINU.BAN | 150 | 20 | 1 | 30 | 100 | No | Serotiny | 0 | 25 | boreal | Pioneer |
| *Pinus resinosa* | PINU.RES | 200 | 40 | 2 | 12 | 275 | No | Seeding | 1 | 15 | boreal | Pioneer |
| *Pinus strobus* | PINU.STR | 300 | 20 | 3 | 100 | 250 | No | Seeding | 1 | 15 | Therm. | M/L- succ. |
| *Populus tremuloides* | POPU.TRE | 150 | 20 | 1 | 1000 | 5000 | Yes | Resprout | 0 | 25 | boreal | Pioneer |
| *Quercus rubra* | QUER.RUB | 250 | 30 | 3 | 30 | 3000 | Yes | Resprout | 1 | 15 | Therm. | M/L- succ. |
| *Thuja occidentalis* | THUJ.OCC | 300 | 30 | 5 | 45 | 60 | No | Seeding | 1 | 15 | boreal | M/L- succ. |
| *Tsuga canadensis* | TSUG.CAN | 300 | 60 | 5 | 30 | 100 | No | Seeding | 1 | 15 | Therm. | M/L- succ. |

\* Thermophilous (Therm.): $\text{minGDD} \geq 500$ AND $\text{maxGDD} \geq 4000$ (see table S2). All other species were considered boreal
\*\* Pioneer: $\text{Shade tolerance} \leq 2$ and $\text{Longevity} \leq 200$ years. All other species were considered mid- to late-successional (M/L – succ.)
† Index of the ability of species to establish under varying light levels where 1 is the least shade tolerant and 5 is the most shade tolerant.
‡ Distance within which 95 % of seeds disperse.

**Table S2.** Select input parameters specific to PICUS for species simulated within the study area.

| Species | Soil nitrogen* | Minimum soil<br>pH† | Maximum soil<br>pH† | Minimum GDD<br>(Base temp 5°C)<br>‡‡ | Maximum GDD<br>(Base temp 5°C)<br>‡‡ | Maximum SMI§ | Optimum SMI§ |
| --- | --- | --- | --- | --- | --- | --- | --- |
| ABIE.BAL | 2 | 2 | 9 | 150 | 2723 | 0.3 | 0 |
| ACER.RUB | 2 | 2 | 9.5 | 500 | 6608 | 0.5 | 0.05 |
| ACER.SAH | 2 | 1.7 | 9.9 | 450 | 5093 | 0.3 | 0 |
| BETU.ALL | 2 | 2 | 10 | 500 | 4517 | 0.5 | 0.05 |
| BETU.PAP | 2 | 2.2 | 9.4 | 150 | 3081 | 0.5 | 0.05 |
| FAGU.GRA | 2 | 2.1 | 9 | 500 | 5602 | 0.7 | 0.1 |
| LARI.LAR | 1 | 3 | 9.6 | 150 | 2548 | 0.3 | 0 |
| PICE.GLA | 3 | 2 | 10.2 | 150 | 2495 | 0.5 | 0.05 |
| PICE.MAR | 2 | 2 | 8.5 | 150 | 2495 | 0.3 | 0 |
| PICE.RUB | 2 | 2 | 7.8 | 450 | 3239 | 0.3 | 0 |
| PINU.BAN | 1 | 2.5 | 10.2 | 300 | 3188 | 0.7 | 0.1 |
| PINU.RES | 1 | 2.5 | 8 | 500 | 3300 | 0.7 | 0.1 |
| PINU.STR | 2 | 2 | 9.3 | 500 | 4261 | 0.7 | 0.1 |
| QUER.RUB | 2 | 2.3 | 11 | 150 | 3024 | 0.5 | 0.05 |
| POPU.TRE | 2 | 2.3 | 11 | 150 | 3024 | 0.5 | 0.05 |
| THUJ.OCC | 2 | 3 | 10 | 500 | 3383 | 0.7 | 0.1 |
| TSUG.CAN | 2 | 2.2 | 9 | 500 | 4660 | 0.5 | 0.05 |

\* Nitrogen response curves: Three classes (1-3) with 1 being very tolerant
† USDA plant fact sheets (USDA, 2016) and the Ontario Silvics Manual (OMNR, 2000) were used to derive the widest optimum pH range possible.
‡ Growing Degree Days (GDD). We used McKenney et al. (2011) growing season model, specifically the minimum GDD for the 0°C and growing season window with degree days over 5°C. For the maximum GDD, we used GDD Maximum from McKenney's previous growing season model (McKenney et al. 2007).
§ Soil Moisture Index (SMI). Determines each species tolerance to drought (see Lexer and Honninger pg. 52). HighTolerance (0.1 to 0.7), MedTolerance (0.05 to 0.5 ), LowTolerance (0 to 0.3).

**Supplementary Material S2.**
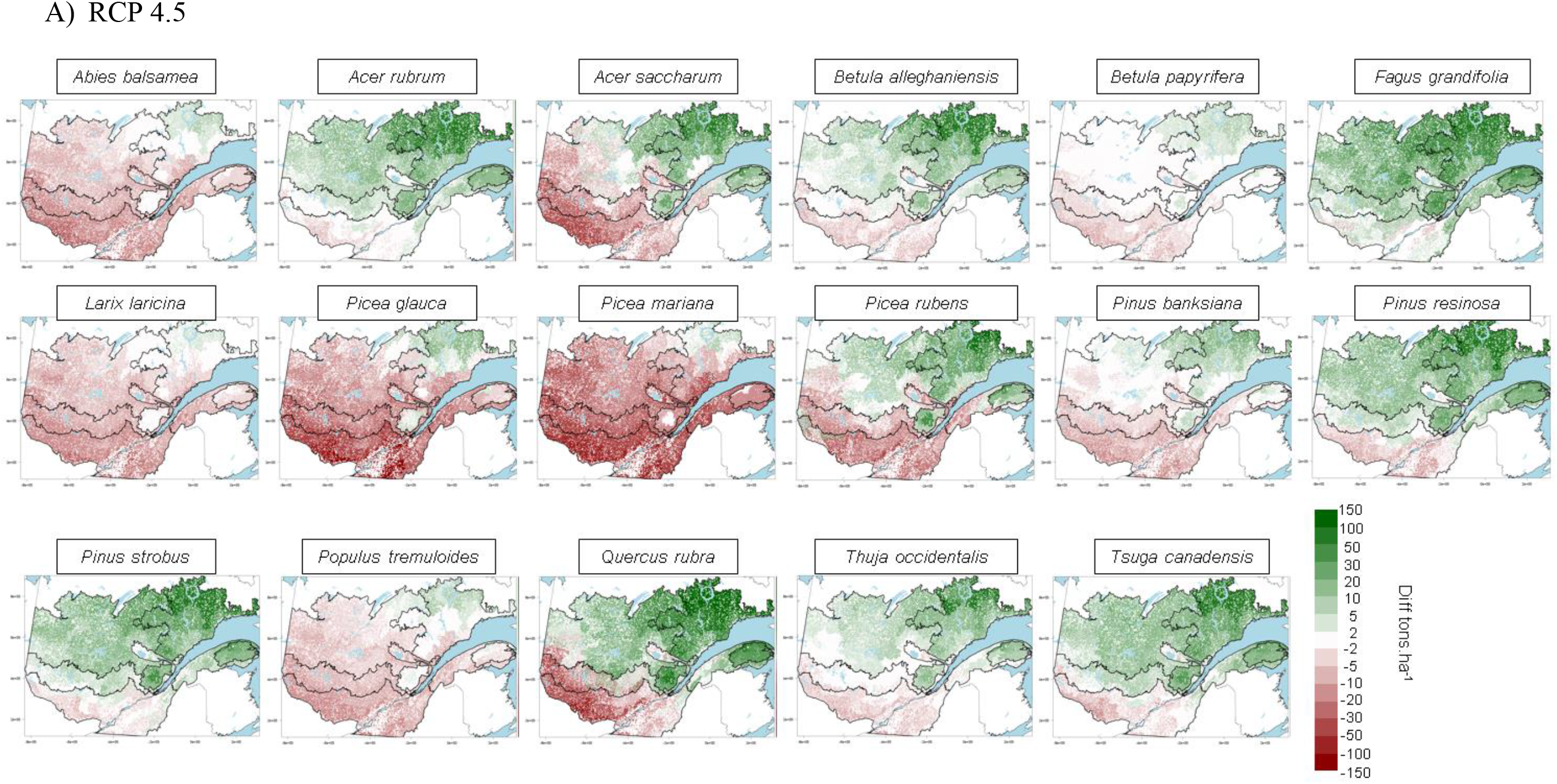
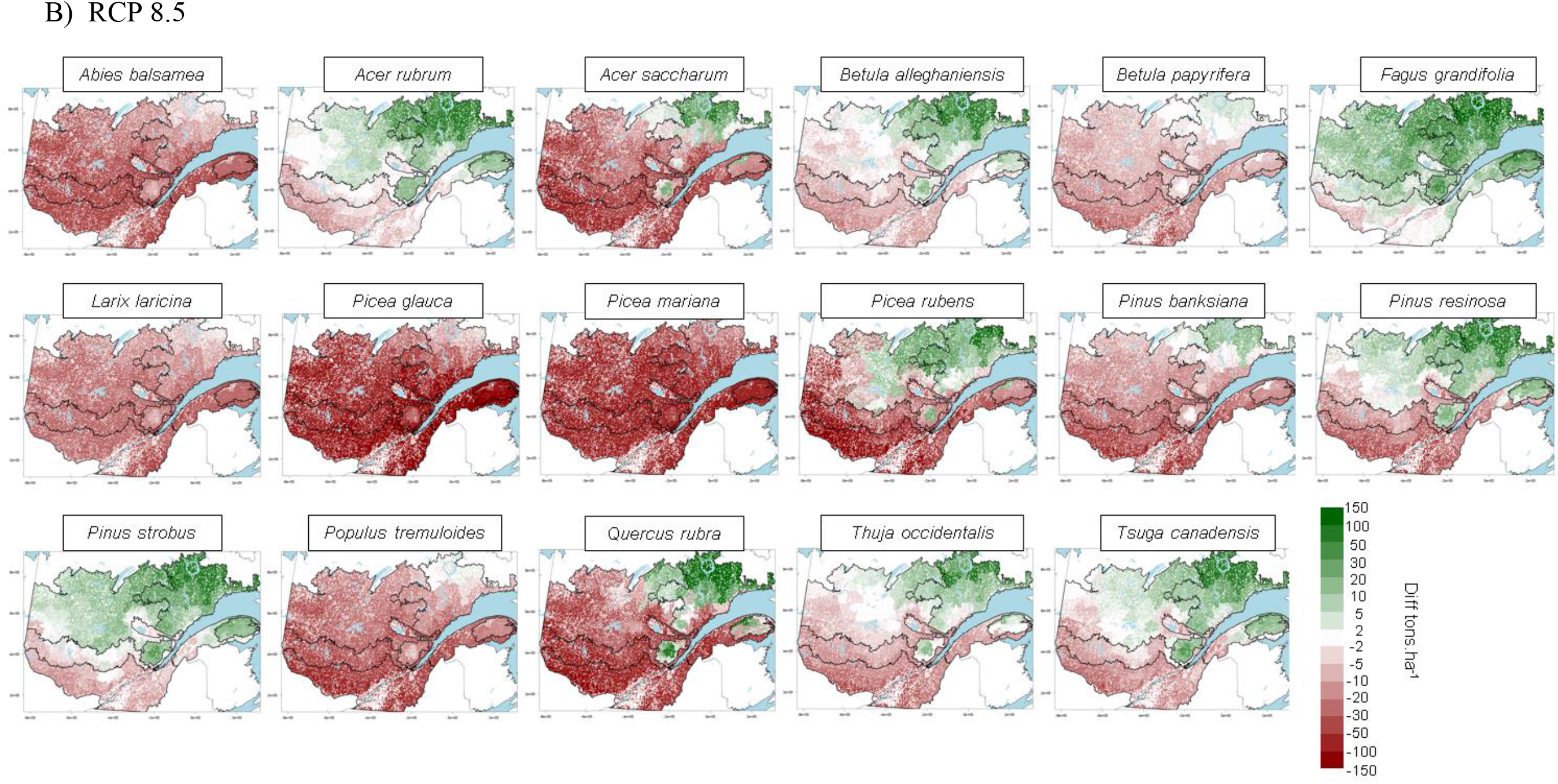
Maps showing for each tree species the difference in landtype-level *maxAGB* (tons per ha) as assessed from PICUS under A) RCP 4.5 or B) RCP 8.5 for the 2071-2100 period with the one simulated under historical climate. Forest regions are outlined in black. *maxAGB* was one of the dynamic inputs used in the Biomass Succession extension in LANDIS-II. We only show *maxAGB* since other dynamic inputs (*SEP* and *maxANPP*) were highly correlated to *maxAGB* (data not shown). See the Material and Methods section for details regarding the calculation of *maxAGB*.

**Supplementary Material S3.**
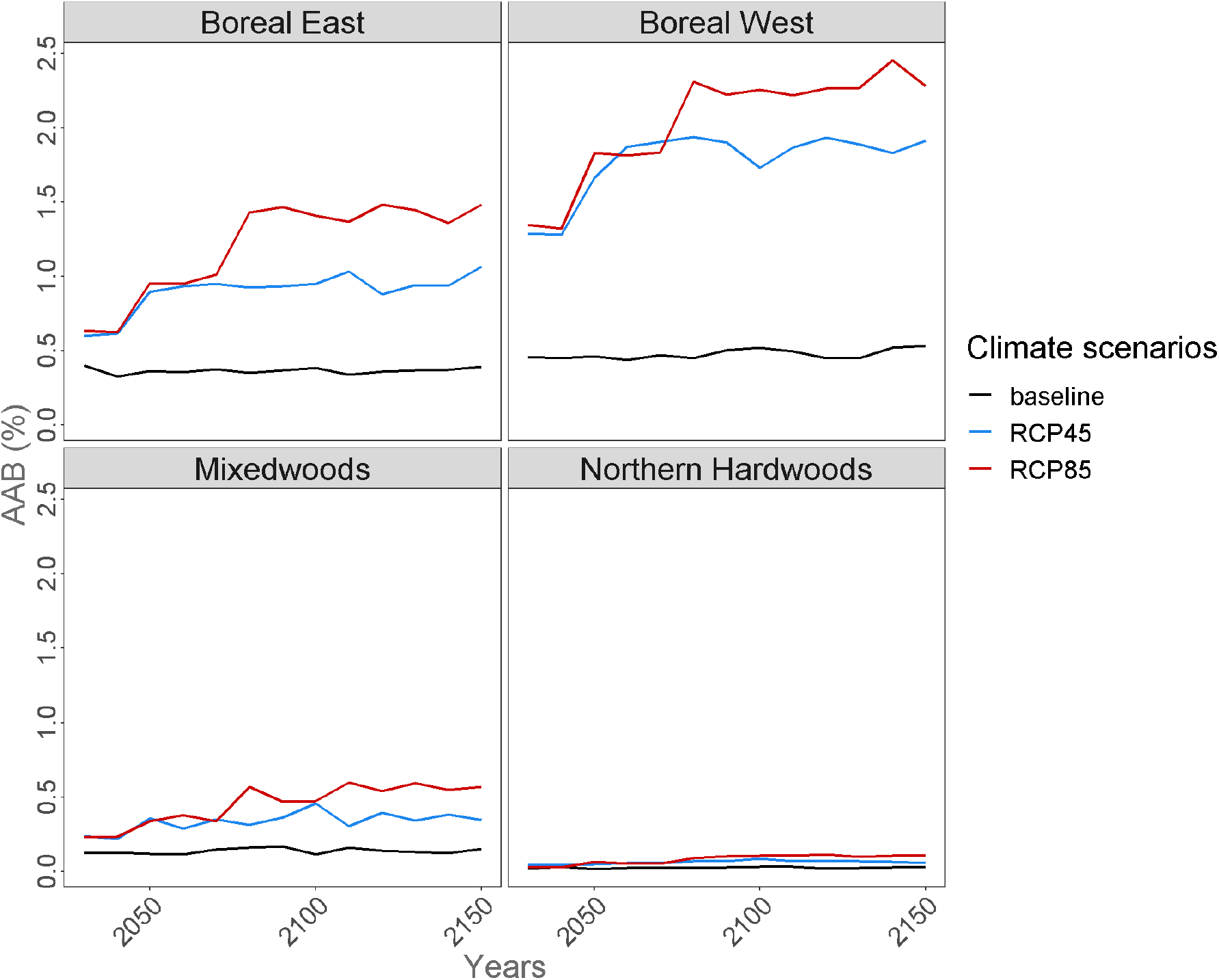
Annual area burned (%) simulated within each of the four forest regions under baseline, RCP 4.5 and RCP 8.5 climate respectively. Values are only showed for simulations including EBFM harvest and are averaged across the five replicates

**Supplementary Material S4.**
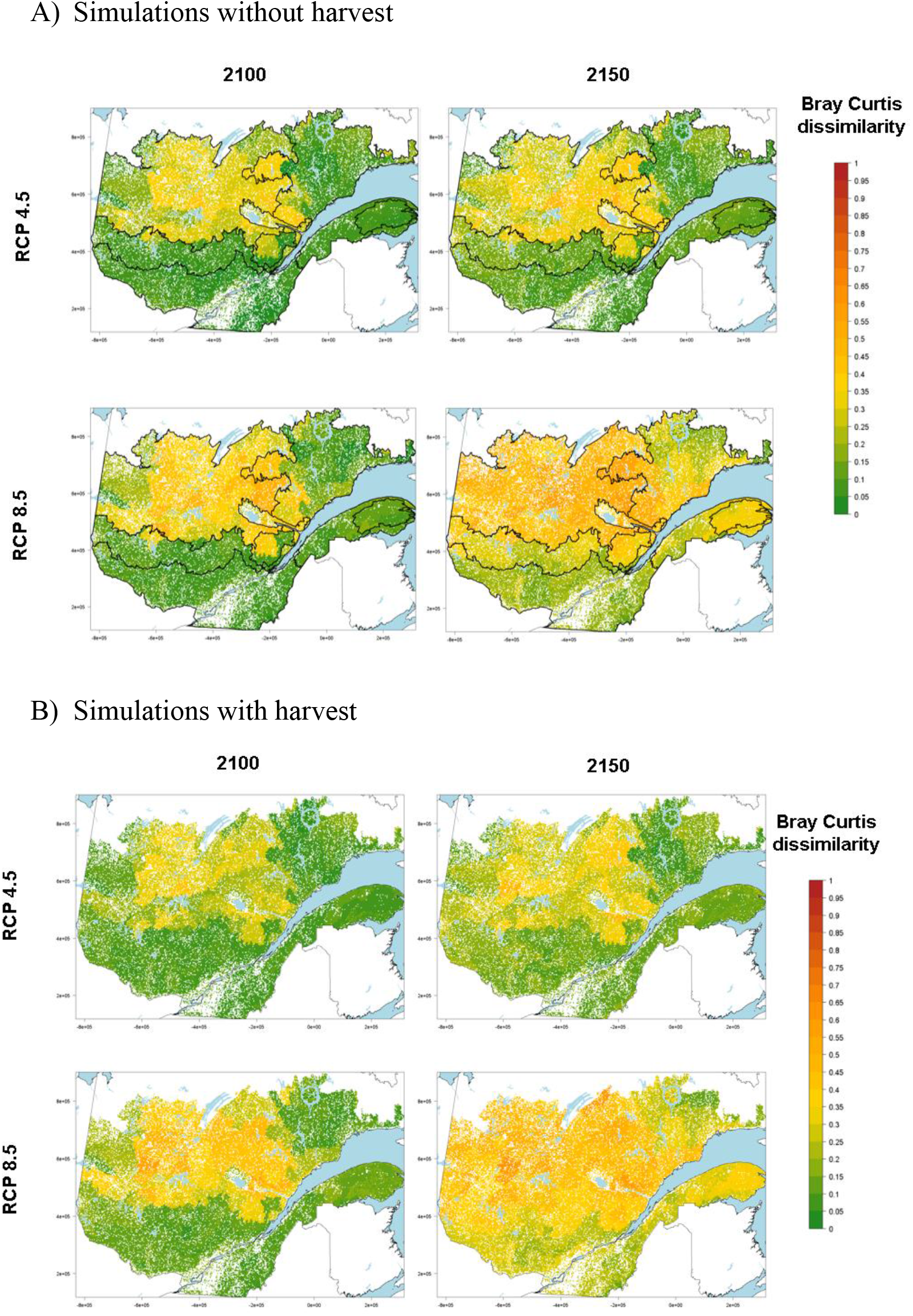
Maps showing Bray-Curtis dissimilarity at the landtype scale. Results are shown at years 2100 and 2150 for simulations ran under the RCP 4.5 and RCP 8.5 climate forcing separately. Forest regions are outlines in black.

**Supplementary Material S5.**
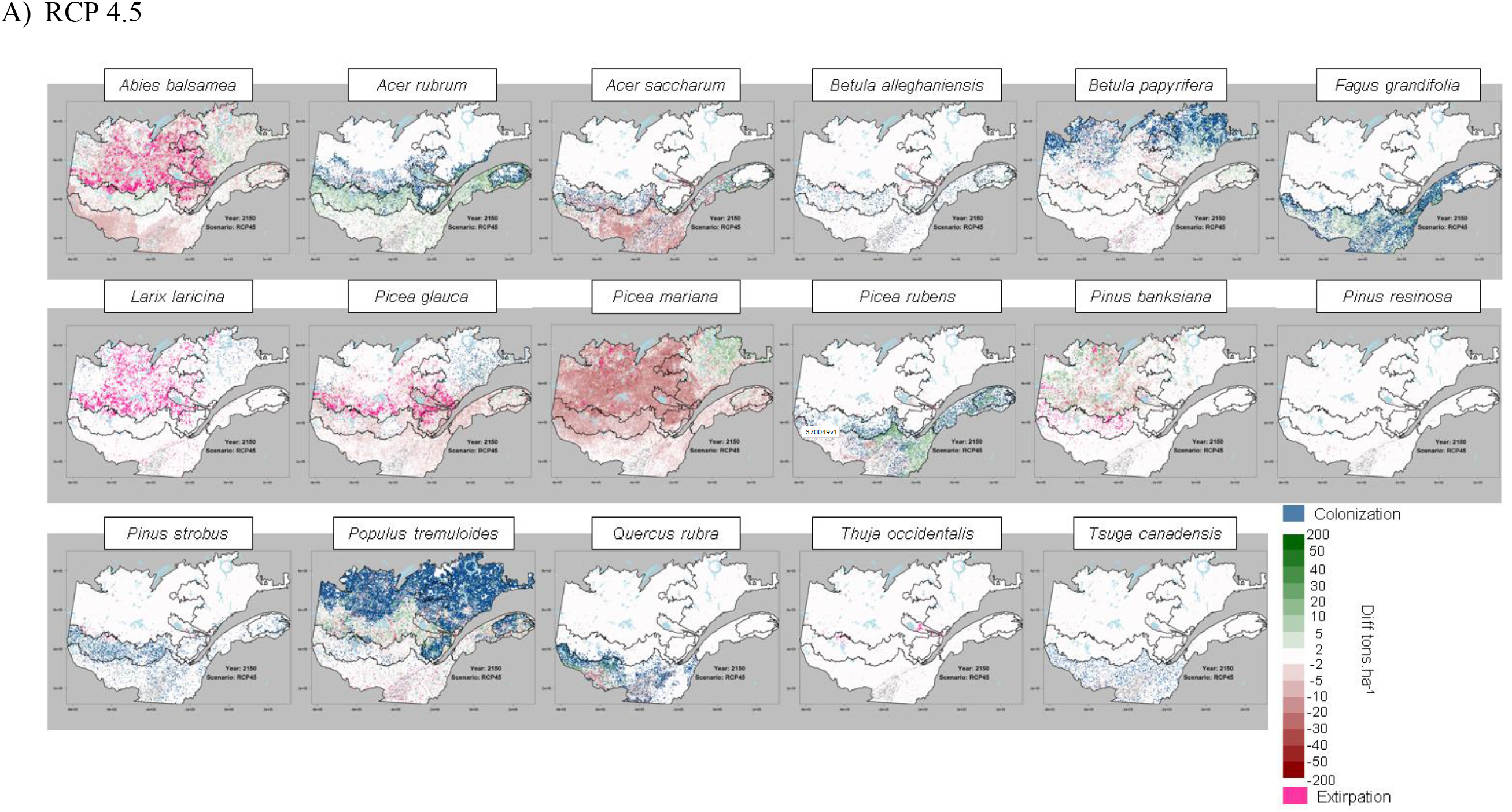
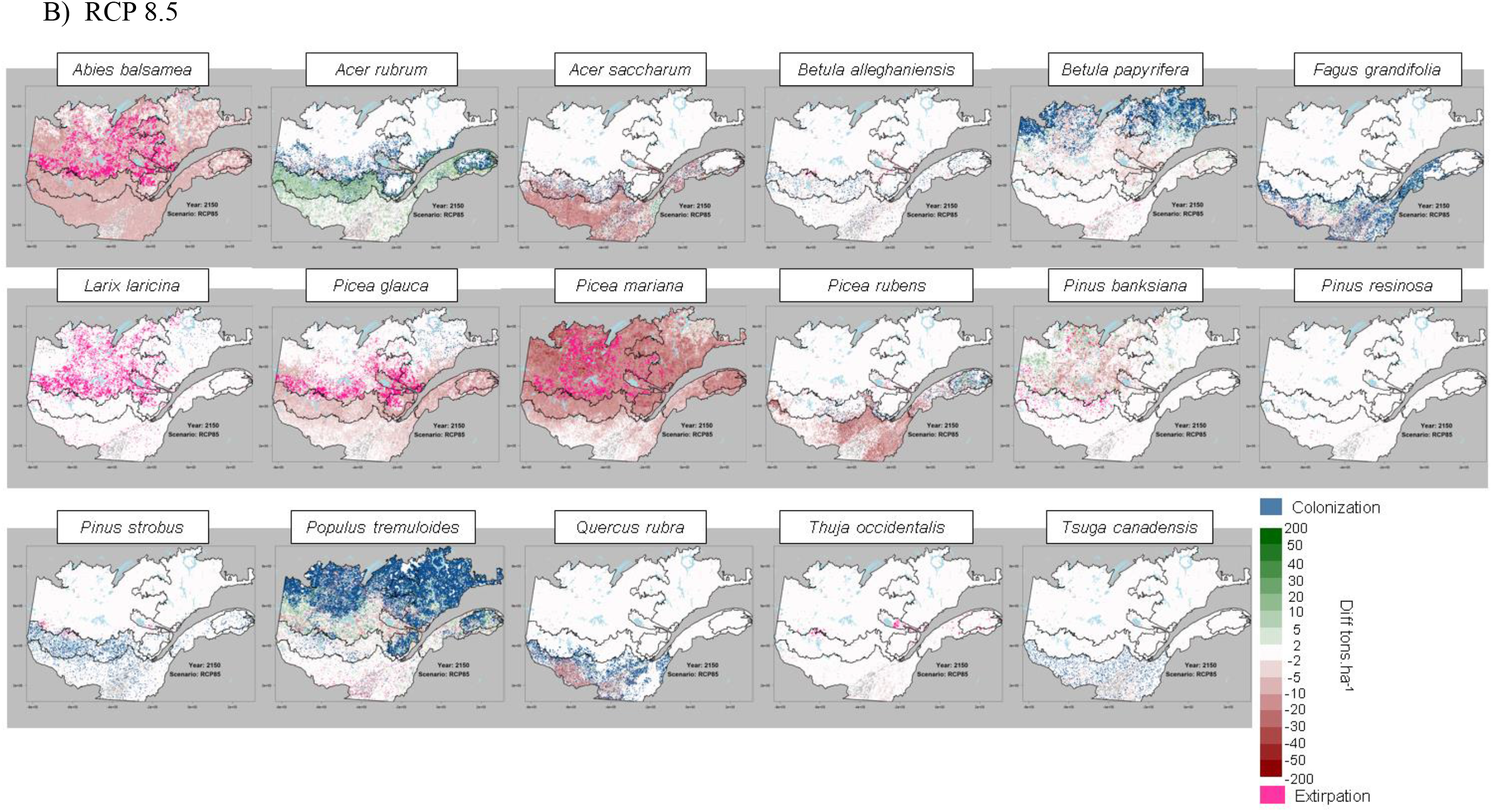
Maps showing for each tree species the difference in AGB (tons per ha) under A) RCP 4.5 or B) RCP 8.5 in 2150 with the one simulated under historical climate in 2150. We also show pixels where species either colonized (blue) or were extirpated (pink). To be extirpated, a species AGB in 2020 must be higher than its 1^st^ percentile (as calculated from the 2020 maps) and drop to nil in 2150. For a pixel to be colonized, the species must be absent (AGB = 0) in 2020 and must be higher than its 1^st^ percentile (as calculated from the 2020 maps) in 2150. We are only showing results where EBFM harvest was included. Forest regions are outlined in black.

**Supplementary Material S6.**
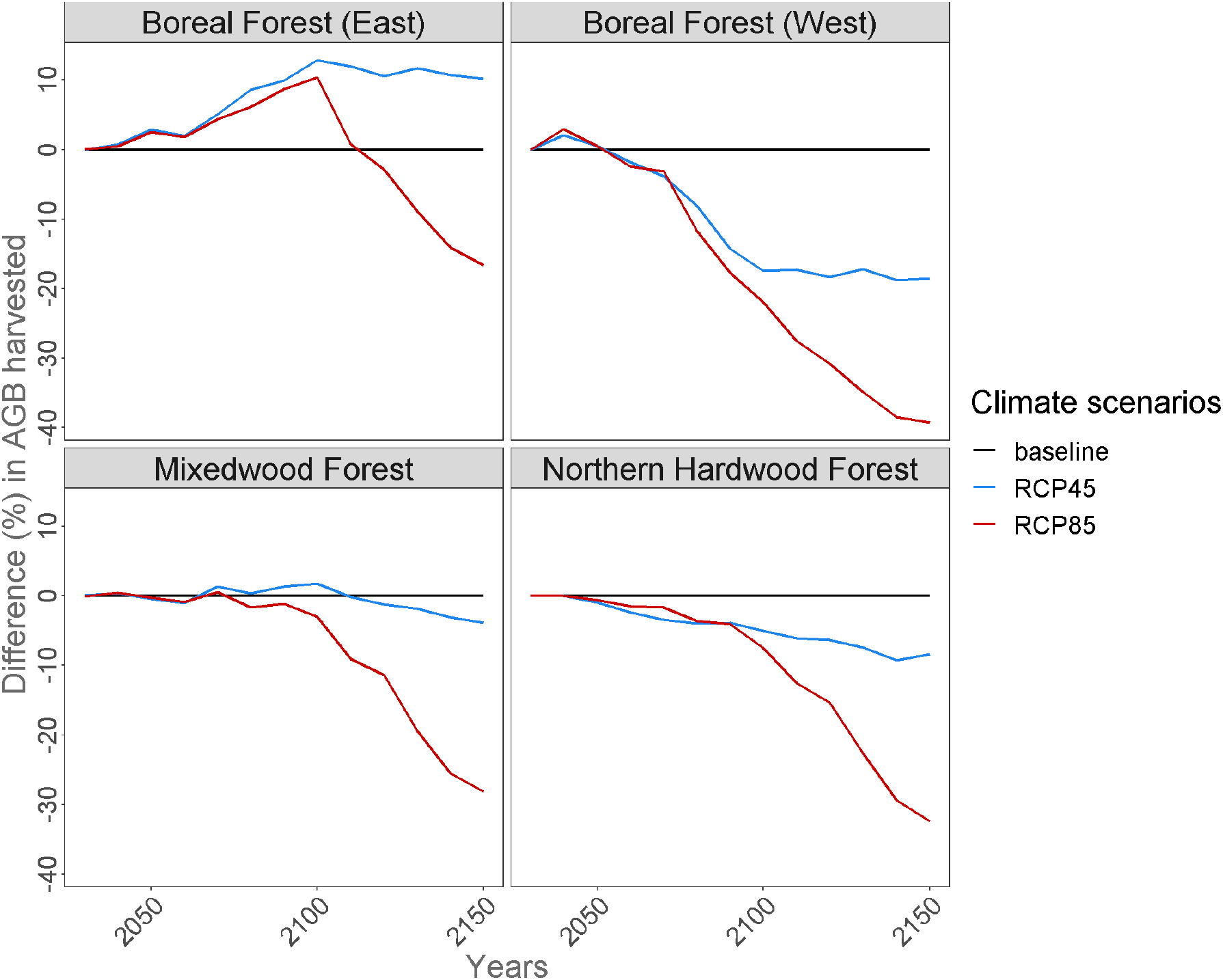
Temporal trends in AGB harvested under the three climate and two harvest scenarios. Results are expressed as differences (%) with AGB harvested at time *t* under baseline climate.

